# Pharmacologic decoupling of IRBC activation from anabolic collapse redefines ribosome biogenesis inhibition as a selective tumor suppressive strategy

**DOI:** 10.64898/2026.08.24.744089

**Authors:** Sandra Menoyo, Berta Forcada, Zoi Mastora, Pau Bosch-i-Crespo, Francisco D. Morón-Duran, Cristina Santos, Ramón Salazar, Antonio Gentilella

## Abstract

Ribosome biogenesis (Ri-Bi) is widely targeted in cancer therapy, yet its inhibition is generally viewed as a broadly anti-anabolic intervention. In colorectal cancer, frontline treatments such as FOLFOX partly disrupt Ri-Bi, eliciting two biologically distinct outputs: an early p53-dependent checkpoint activation, known as the impaired ribosome biogenesis checkpoint (IRBC), and a later global anti-anabolic collapse associated with toxicity and limited durability. At clinically relevant doses, these outputs have been considered pharmacologically inseparable. Here we demonstrate that Ri-Bi inhibition can be functionally dissociated and selectively tuned toward checkpoint engagement. Using a genome-engineered Venus-RPL11 reporter and *TP53* isogenic colorectal cancer models, we show that combining sub-effective doses of mechanistically distinct Ri-Bi inhibitors reprograms the cellular response toward dominant IRBC-mediated p53 activation while minimizing p53-independent cytotoxicity. This dose architecture induces profound growth suppression exclusively in *TP53*-proficient cells and prevents adaptive outgrowth during prolonged treatment. Importantly, pharmacologic rescue of mutant p53 (R175H) with arsenic trioxide restores IRBC responsiveness, extending this framework to genetically advanced disease. Together, our findings establish that ribosome biogenesis inhibition can be selectively directed toward nucleolar surveillance activation, redefining Ri-Bi targeting as a checkpoint-based therapeutic principle.

## Introduction

Ribosomes are the molecular machines responsible for protein synthesis and are therefore essential for virtually all cellular processes. Their production, known as ribosome biogenesis (Ri-Bi), is an energetically demanding and tightly coordinated pathway that couples cellular growth to biosynthetic capacity [1, 2]. In cancer, increased proliferative requirements drive sustained upregulation of Ri-Bi, making it a widespread metabolic hallmark of malignant cells [3–5].

To preserve cellular homeostasis, eukaryotic cells have evolved a surveillance mechanism that monitors the integrity of ribosome biogenesis [6, 7]. Perturbations in Ri-Bi activate a nucleolar stress response—referred to as the impaired ribosome biogenesis checkpoint (IRBC)—that stabilizes p53 and enforces anti-proliferative outcomes [8–11]. Mechanistically, IRBC activation is mediated by a ribonucleoprotein complex composed of ribosomal proteins L5 (uL18), L11 (uL5), and 5S rRNA. Under normal conditions, this complex is incorporated into the pre-60S ribosomal subunit, whereas upon ribosome biogenesis impairment it is redirected to inhibit the E3 ubiquitin ligase MDM2, thereby preventing p53 degradation and enabling rapid p53 stabilization [9–12]. This dual role positions the L5-L11-5S complex as pivotal relay activating cellular anabolism or tumor suppression, depending on the biological context. Hallmark IRBC dynamics include nucleolar recruitment/retention of L11/L5 and assembly of the 5S-RNP–MDM2 complex [6, 9, 13]. Consistently, disruption of the MDM2–5S-RNP interaction abolishes p53 activation by ribosomal stress while preserving DNA damage responses, and in vivo abrogation of this pathway accelerates MYC-driven tumorigenesis [14, 15].

Importantly, perturbation of Ri-Bi produces two biologically distinct outputs: an early activation of the IRBC tumor suppressive pathway and a later global reduction in ribosome production that limits cellular anabolic capacity. While the former represents a selective checkpoint response, the latter reflects a broader collapse of protein synthesis that contributes to non-specific cytotoxicity [1, 14].

From a pharmacological perspective, these outputs are largely inseparable at clinically relevant doses. Several agents used in colorectal cancer (CRC) treatment perturb Ri-Bi at distinct steps of the rRNA life cycle. The antimetabolite 5-fluorouracil (5-FU), beyond thymidylate synthase inhibition, is incorporated into rRNA and disrupts pre-rRNA processing [16, 17]. Oxaliplatin induces p53 activation through ribosome biogenesis stress and disrupts nucleolar organization [18, 19]. Actinomycin D directly blocks rDNA transcription and remains a prototypical inducer of nucleolar stress [1, 10]. These agents therefore provide complementary entry points to pharmacologically perturb Ri-Bi, especially in CRC.

Clinically, the therapeutic potential of Ri-Bi inhibition is constrained by dose-dependent toxicities and by the emergence of resistance in advanced disease [20], likely reflecting the dominant contribution of global anabolic collapse at high drug concentrations. This has prompted interest in strategies that preserve tumor suppressive signaling while minimizing non-specific cytotoxicity.

One emerging approach is “multiple low-dose” (MLD) therapy, in which partial inhibition of several nodes within a pathway collectively produces strong biological effects while reducing toxicity and delaying resistance. In EGFR-mutant lung cancer models, low-dose multi-target inhibition of the MAPK pathway effectively suppresses signaling and prevents adaptive resistance while remaining well tolerated in vivo [21]. We reasoned that a similar strategy could be applied to ribosome biogenesis by combining sub-effective doses of agents targeting distinct steps of the rRNA life cycle together with pharmacological amplification of p53 signaling, thereby selectively engaging the IRBC while limiting global anabolic collapse.

To test this, we developed a genome-engineered Venus-L11 reporter that enables dynamic visualization of IRBC activation in living colorectal cancer cells. Using this platform together with *TP53* isogenic models, we show that coordinated multiple low-dose targeting of Ri-Bi, combined with the MDM2 antagonist CGM-097 [22, 23], selectively amplifies IRBC-dependent p53 activation, resulting in robust and durable growth suppression in *TP53*-proficient cells while sparing *TP53*-null counterparts. Furthermore, pharmacological reactivation of mutant p53 with arsenic trioxide (ATO) restores IRBC responsiveness in cells harboring the R175H mutation, extending this strategy to advanced CRC [24–26]. Together, these findings demonstrate that IRBC activation can be pharmacologically decoupled from global anabolic collapse and establish selective checkpoint engagement as a strategy to exploit ribosome biogenesis in cancer.

## Results

### Venus-RPL11 is a robust tool for studying IRBC activation

With the aim of better understanding the dynamics of the IRBC response as a function of ribosome biogenesis stress, we generated a CRISPR-Cas9 knock-in of Venus at the endogenous locus of ribosomal protein L11 (RPL11), a core component of the IRBC complex (Fig. 1a). The Venus tag was fused in frame to the N-terminus of L11, which is exposed on the ribosomal outer shell (**Fig. S1a**), predicting minimal interference with ribosome biogenesis and function. Venus-positive cells were FACS sorted, clonally expanded and further characterized. Western blot analysis confirmed the expression of a ∼45 kDa Venus-L11 fusion protein detected by both anti-GFP and anti-L11 antibodies, alongside the ∼20 kDa endogenous untagged L11 (**Fig. 1b**). To assess whether the tagged protein retained normal ribosomal function, we performed polysome profiling by sucrose gradient ultracentrifugation. Venus fluorescence and rRNA content were monitored across fractions. Venus-L11 co-sedimented with 60S subunits, monosomes and polysomes, paralleling endogenous L11 distribution (**Fig. 1c**), indicating that the tagged protein is efficiently incorporated into translationally competent ribosomes.

**FIGURE 1.**
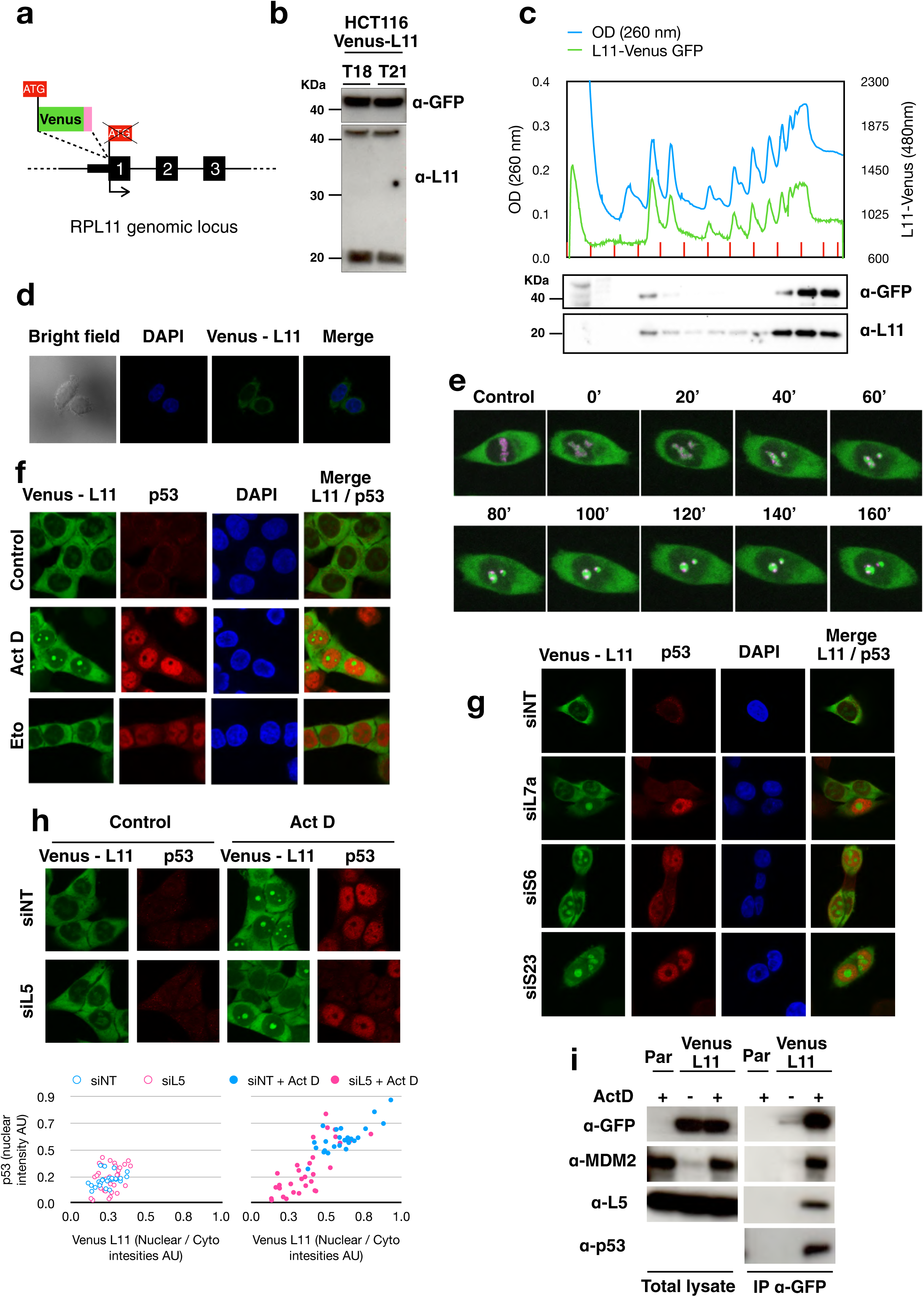
Venus-L11 is a dynamic and specific reporter of IRBC activation. (a) Schematic representation of CRISPR-Cas9–mediated knock-in of Venus in frame with the N-terminus of endogenous RPL11. (b) Western blot analysis of HCT116 Venus-L11 monoclonal cultures (T18, T21) showing expression of the ∼45 kDa Venus-L11 fusion protein detected by anti-GFP and anti-L11 antibodies. Untagged L11 (∼20 kDa) is detected in knock-in cells. (c) Polysome profiling of HCT116 Venus-L11 (T18) cells by sucrose gradient ultracentrifugation. Venus fluorescence (480 nm) and rRNA absorbance (260 nm) were monitored across the grandient. *Bottom panel*: western blot of methanol-chloroform precipitated proteins from corresponding sucrose gradient fraction. anti-GFP and anti-L11 antibodies were used to detect both tagged and untagged L11 proteins.. (d) Live-cell imaging of HCT116 Venus-L11 cells under basal growing conditions. (e) Time-lapse microscopy following Actinomycin D treatment in HCT116 Venus-L11 cells transfected with mPlum-FBL. Venus-L11 progressively accumulates in Fibrillarin (FBL)-positive nucleolar structures within 160 min. (f) Immunofluorescence of HCT116 Venus-L11 cells upon a ribosomal stress (Actinomycin D) and a genotoxic stress (Etoposide). Immunodetection of p53 with an anti-p53 antibody and Venus signal were detected. DAPI stained nuclei. (g) Knockdown of ribosomal proteins S6, S23 and L7a ribosomal proteins induces nucleolar accumulation of Venus-L11 and correlates with p53 stabilization, as revealed by immunofluorescence. (h) Representative immunofluorescence images of acute L5 knockdown in basal condition (Control) and under Actinomycin D (Act D). Venus-L11 and p53 levels were detected. Quantification of nuclear/cytoplasmic Venus signal and of nuclear p53 intensities are shown. (i) Anti-GFP immunoprecipitation of post-ribosomal lysates from HCT116 Venus-L11 cells in control conditions and following Actinomycin D treatment. To detect IRBC complex engagement Immunoblotting revealed L5, MDM2 p53. Cell lysates from Parental HCT116 cells treated with Actinomycin D were used as negative control of immunoprecipitation.

Live-cell imaging under basal conditions showed that Venus-L11 is predominantly cytoplasmic, consistent with incorporation into mature ribosomes, while a smaller nuclear fraction localizes to nucleolar structures, as confirmed by co-localization with the nucleolar marker Fibrillarin (**Fig. 1d-e**). Upon ribosome biogenesis stress induced by Actinomycin D, Venus-L11 rapidly accumulated in Fibrillarin-positive nucleolar compartments within ∼160 minutes (**Fig. 1e**), accompanied by a modest increase in diffuse nucleoplasmic signal.

To validate Venus-L11 as a specific reporter of IRBC activation, we compared Actinomycin D a ribosomal stress inducer [10], with Etoposide, a genotoxic stressor [27]. While both treatments induced robust nuclear p53 accumulation, nucleolar relocalization of Venus-L11 occurred exclusively in Actinomycin D–treated cells (**Fig. 1f**), demonstrating that L11 redistribution is specific to ribosome biogenesis stress and constitutes a real-time marker of IRBC activation, in agreement with previous observations [9, 28]. Consistently, acute suboptimal knockdown of L5, a core component of the IRBC complex (**Fig. 2b**), largely impaired Venus-L11 nucleolar accumulation in those cells lacking p53 stabilization under Actinomycin D treatment (**Fig. 1h**), indicating that L11 relocalization depends on intact IRBC complex assembly. Conversely, knockdown of ribosomal proteins of the 40S (S6, S23) or 60S (L7a) subunits—previously shown to activate IRBC signaling [8, 10, 29, 30]—induced robust nucleolar accumulation of Venus-L11 that correlated with p53 stabilization (**Fig. 1g**). Finally, anti-GFP pulldown of Venus-L11 following Actinomycin D treatment co-immunoprecipitated L5, MDM2 and p53 (**Fig. 1i**). Collectively, these results establish Venus-L11 as a faithful and dynamic reporter of IRBC activation, thus providing a unique standpoint to study the IRBC signaling and to evaluate therapeutic strategies targeting ribosome biogenesis.

**FIGURE 2.**
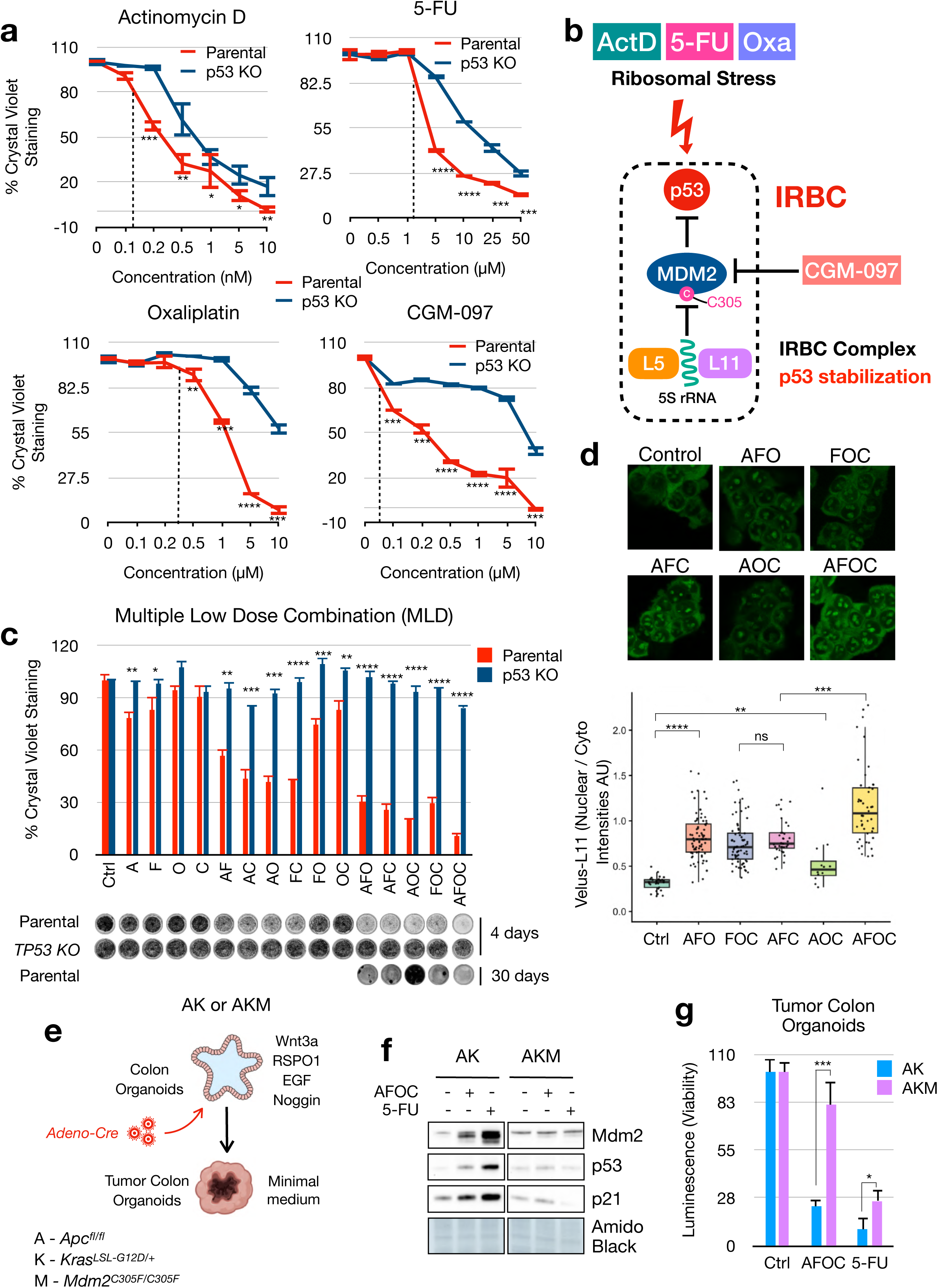
Multiple low-dose regimens selectively engage the IRBC through the 5S-RNP–MDM2–p53 axis. (a) Dose–response curves of parental and TP53-knockout HCT116 cells treated for 4 days with Actinomycin D, 5-FU, Oxaliplatin, or CGM-097. Cell viability was assessed by crystal violet staining and expressed relative to untreated controls (set as 100% viability). Data represent mean ± SD from three biological replicates. Statistical comparisons were performed between parental and TP53-knockout cells at each concentration. (b) Schematic representation of IRBC-mediated p53 stabilization via the L5/L11/5S rRNA–MDM2 axis. (c) Multiple low-dose (MLD) combinatorial treatments in parental and TP53-knockout HCT116 cells. Cells were treated for 4 days with single, double, triple, or quadruple combinations of Actinomycin D (A), 5-FU (F), Oxaliplatin (O), and CGM-097 (C) at IC10–15 concentrations. Cell growth was assessed by crystal violet staining and expressed relative to untreated controls. Bars represent mean ± SD from three biological replicates. Statistical comparisons were performed between parental and TP53-knockout cells within each treatment condition. Representative stained wells are shown below. Chronic exposure (30 days) revealed adaptive outgrowth in all regimens except the quadruple AFOC combination. (d) Representative fluorescence images of HCT116 Venus-L11 distribution in response to MLD combinations. Nuclear/cytoplasmic ratios were quantified and plotted for every condition (bottom panel). Statistical comparisons were performed between each MLD condition and untreated control using two-tailed unpaired Student’s *t*-test. (e) Schematic representation of tumor colon organoid generation. Colon organoids derived from AK and AKM mice were infected with Adeno-Cre to induce *Apc* deletion and *Kras*^G12D^ activation, and subsequently selected in minimal medium to establish tumor colon organoids. (f) AK and AKM tumor colon organoids were treated for 24 hours with AFOC or 10 μM 5-FU. Whole-cell lysates were analyzed by western blot for Mdm2, p53 and p21 expression. Amido Black staining is shown as loading control. (g) AK and AKM tumor colon organoids were treated for 4 days with AFOC or 10 μM 5-FU. Cell viability was measured using CellTiter-Glo and expressed relative to untreated controls. Bars represent mean ± SD from six replicate wells per condition. Statistical significance was assessed by comparing AK and AKM organoids within each treatment condition using two-tailed unpaired Student’s *t*-test.

### Assessing the tumor suppressive potential of IRBC in colorectal cancer cells

We reasoned that selectively activating the p53-dependent tumor suppressive arm of the IRBC could reinforce the anti-proliferative response to ribosome biogenesis perturbation, particularly in colorectal cancer, where FOLFOX remains a first-line therapy. Specifically, we aimed to discriminate between the global anti-proliferative effects of ribosome biogenesis inhibition— stemming from reduced ribosome production and anabolic collapse—and the specific p53-dependent tumor suppressive output mediated by IRBC activation.

Dose titration of Actinomycin D in HCT116 cells showed that siRNA-mediated knockdown of p53 partially desensitized cells, particularly at intermediate concentrations (**Fig. S1b**), confirming the contribution of p53 to the anti-proliferative response induced by ribosome biogenesis stress. To rigorously define the amplitude of the IRBC response, we sought to use as a reference a total *TP53* KO genetic setting. Notably, the widely used HCT116 p53^−/−^ line obtained targeting the exon 2 of *TP53* [31] remained as sensitive to Actinomycin D as parental cells (**Fig. S1c**), potentially due to N-terminally truncated p53 isoforms [32]. Hence, we generated an isogenic *TP53* knockout by targeting exon 5, which is shared by all annotated TP53 transcripts (**Fig. S1d**). Clone #8, which lacked detectable p53 even after Actinomycin D treatment, was selected for subsequent experiments (**Fig. S1e**).

Dose-response assays in parental and *TP53* KO cells demonstrated that *TP53*-proficient cells are significantly more sensitive to Actinomycin D, 5-FU, Oxaliplatin or the MDM2 inhibitor CGM-097 (**Fig. 2a**), consistent with p53-mediated tumor suppressive activity. Western blot analysis confirmed selective stabilization of p53 and induction of its transcriptional targets p21 and MDM2 exclusively in parental cells (**Fig. S2a-d**). With this approach we identified sub-effective drug concentrations producing approximately 10–15% reduction in viability (IC10–15), hereafter referred to as low dose (LD), which had minimal impact on *TP53* KO cells. These conditions allowed preferential engagement of p53-dependent responses while limiting p53-independent cytotoxicity, which in the case of Act-D, 5-FU and Oxaliplatin reflects the activation of the IRBC response. Using these LD conditions, we tested combinatorial multiple low-dose (MLD) treatments with Actinomycin D (A), 5-FU (F), Oxaliplatin (O) and CGM-097 (C). Most double combinations produced greater growth inhibition than the corresponding single agents in *TP53*-proficient cells, while triple combinations further enhanced the response, as also evidenced by Bliss independence analysis (**Fig. S2e**). Notably, the quadruple combination (AFOC) induced ∼90% inhibition in *TP53*-proficient cells compared to a mere ∼10-15% in *TP53* KO cells (**Fig. 2c** and **Fig. S2f**). Importantly, consistent results were obtained in an independent HCT116 *TP53* KO clone (#3, **Fig. S1e**) (**Fig. S3a**) as well as in a different colorectal cancer cell line (RKO) at its respective IC10-15 dose (**Fig S3b-e**). By contrast, the same regimens in non-transformed WI38 human lung fibroblasts caused only minor growth inhibition (approximately 10–20%; **Fig. S3f**) indicating limited activity in a non-malignant cellular context. Together these findings demonstrate a robust amplification of p53-dependent tumor suppression with minimal p53-independent effects.

Given that resistance to FOLFOX frequently emerges in advanced colorectal cancer, we next examined whether sustained MLD combinations could prevent adaptive outgrowth. Chronic treatment for one month with periodic drug renewal resulted at different extent in resistant clones emergence under all regimens except the quadruple AFOC combination (**Fig. 2c**), suggesting that coordinated low-dose targeting of ribosome biogenesis and p53 signaling limits adaptive resistance. Importantly, consistent with the central role of IRBC activation in mediating this effect, the AFOC MLD regimen induced the strongest nucleolar accumulation of L11 in Venus-L11 HCT116 cells (**Fig. 2d**), supporting a correlation between IRBC activation and therapeutic efficacy.

While validation in an independent CRC cell lines addressed the generalizability of the observations, we reasoned that the most stringent test of our mechanistic model was to determine whether the MLD response specifically depends on the IRBC itself in a *TP53*-proficient model. We therefore extended the analysis to genetically engineered mouse tumor colon organoids (TCOs) derived from Apc^fl/fl^; Kras^LSL-G12D/+^ mice (AK). As an orthogonal genetic approach, we compared these organoids with isogenic TCOs (AKM) carrying the homozygous Mdm2^C305F^ knock-in mutation, which selectively abolishes IRBC-mediated p53 activation while preserving p53 responses to genotoxic stress [14]. We first generated Apc^-/-^; Kras^G12D/+^ and Apc^-/-^; Kras^G12D/+^; Mdm2^C305F/C305F^ TCOs by Adeno-Cre infection followed by minimal medium selection (**Fig. 2e**) and determined the IC10-15 of AK TCOs to the four drugs (**Fig. S4a**). Consistent with the proposed mechanism, the AFOC MLD regimen induced robust p53 activation and growth inhibition in AK organoids but failed to elicit a comparable response in AKM organoids (**Fig. 2f-g**), providing additional genetic evidence that the therapeutic effect of the MLD strategy is mediated specifically through the IRBC pathway rather than through generic p53 activation.

### Multiple Low Dose regimens decouple IRBC activation from Ribosome Biogenesis inhibition

The selective growth inhibition observed in *TP53*-proficient cells suggested that the MLD response was primarily driven by checkpoint activation rather than by a direct loss of biosynthetic capacity. To test this mechanistically, we compared the effects of the AFOC regimen with conventional high-dose ribosome biogenesis inhibition by assessing polysome abundance, global protein synthesis and nascent rRNA synthesis. Polysome profiling showed that, while high-dose 5-FU markedly reduced polysome content in both parental and *TP53* KO HCT116 cells, consistent with a p53-independent impairment of translational capacity. In contrast, AFOC reduced polysome abundance only in parental cells, whereas *TP53* KO cells remained largely comparable to untreated controls (**Fig. 3a,b**). Consistent with these findings, puromycin incorporation revealed that the triple and quadruple MLD regimens caused a partial reduction in global protein synthesis, selectively in *TP53*-proficient cells. By contrast, conventional high-dose Actinomycin D strongly suppressed translation in both parental and *TP53* KO cells (**Fig. 3c**). Importantly, metabolic labelling of nascent RNA with ethynyl uridine showed that AFOC did not detectably reduce rRNA synthesis and maturation in either genotype, opposite to high-dose Actinomycin D which produced a pronounced inhibition of rRNA synthesis (**Fig. 3d**). Together, these results indicate that the MLD regimen does not primarily suppress ribosome production. Rather, the partial reduction in protein synthesis observed in parental cells arises downstream of *TP53*-dependent checkpoint activation, thereby distinguishing selective IRBC engagement from the p53-independent anabolic collapse induced by conventional high-dose treatment.

**FIGURE 3.**
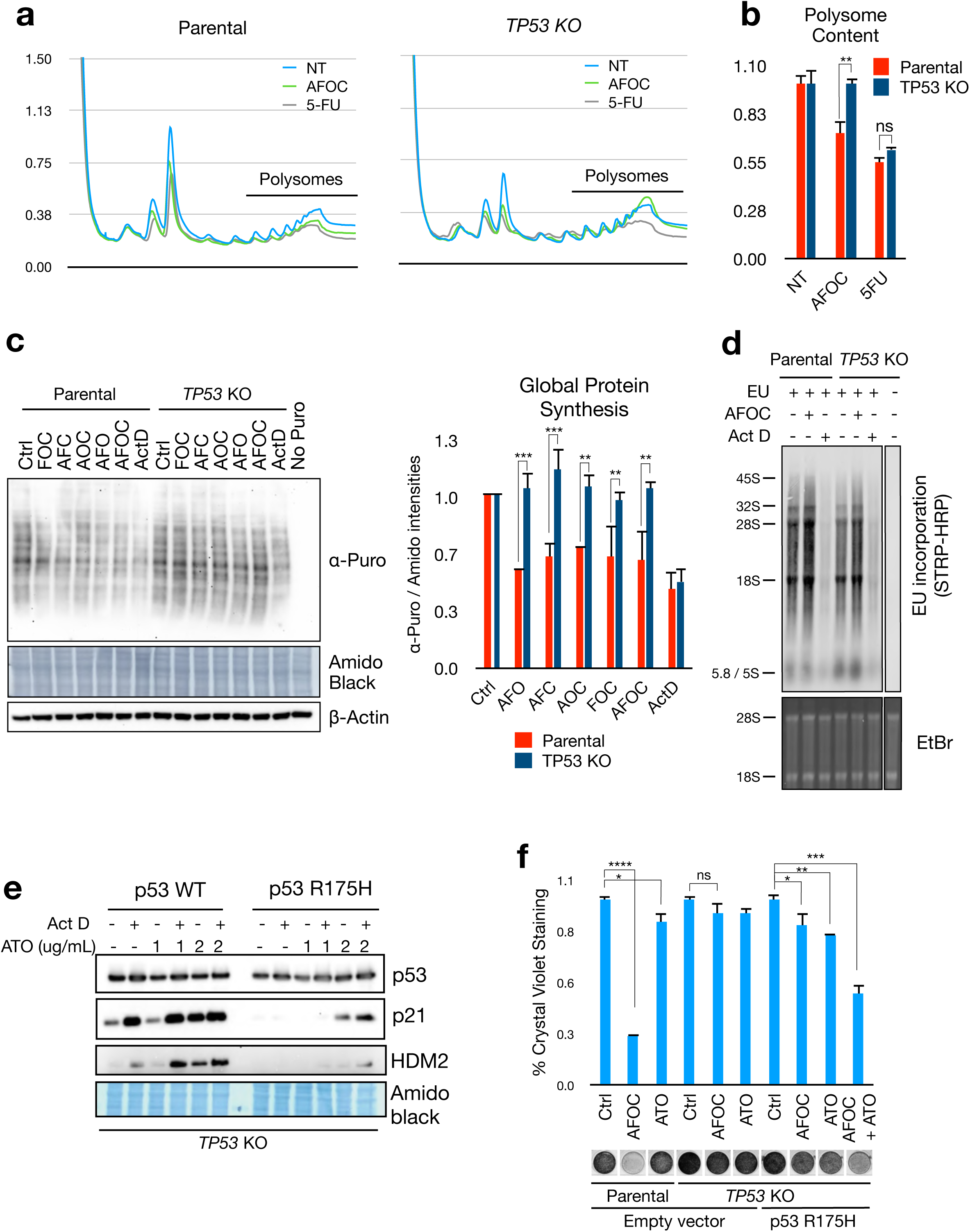
Pharmacological decoupling of IRBC activation from ribosome biogenesis inhibition and anabolic collapse by multiple low-dose regimens. (a) Representative polysome profiles analyses of parental and *TP53* KO HCT116 cells untreated or treated for 24 hours with AFOC or 10 μM 5-FU. The polysomal region used for quantification is indicated. (b) Polysome content was quantified as the area under the curve of the polysomal region and normalized to the corresponding untreated condition. Bars represent mean ± SD from three independent biological experiments. Statistical significance was assessed by comparing parental and *TP53* KO cells within each treatment condition. (c) Parental and *TP53* KO HCT116 cells were treated for 24 hours with the indicated MLD combinations or Actinomycin D (10 nM) and pulsed with puromycin during the final 30 min. Global protein synthesis was assessed by anti-puromycin western blotting. A non-pulsed sample served as negative control. Puromycin signal was normalized to Amido Black staining from the corresponding lane and expressed relative to the untreated control of each genotype. A representative experiment and quantification from three independent biological replicates are shown. (d) Parental and *TP53* KO HCT116 cells were treated for 24 hours with AFOC or Actinomycin D (10 nM) and pulsed with ethynyl uridine during the final 2 hours. Three micrograms of total RNA were resolved by agarose gel electrophoresis, transferred to membrane, conjugated to biotin-azide by click chemistry on the membrane, and nascent RNA was detected with streptavidin–HRP. Ethidium bromide staining is shown as loading control, and cells not pulsed with ethynyl uridine served as negative control. (e) Transient re-expression of wild-type p53 or the R175H mutant in TP53 KO HCT116 cells. Cells were treated Actinomycin D alone or in presence of Arsenic Trioxide (ATO). The p53 transcriptional targets of p21 and MDM2 were detected by western blot. Amido black served as loading control. (f) TP53 KO cells expressing p53 R175H were treated with ATO alone, AFOC (MLD) alone, or ATO + AFOC. Crystal violet assays demonstrate that ATO restores sensitivity of R175H-expressing cells to the AFOC regimen. Western blot confirms reactivation of p53 transcriptional targets upon combined treatment.

### Restoring p53 function expands the therapeutic window of IRBC targeting

Advanced stages of CRC, as well as many other tumor types, frequently harbor hotspot mutations in the TP53 gene that abrogate its tumor suppressive activity [24]. Such mutations represent a major evolutionary bottleneck, enabling tumor progression while simultaneously conferring resistance to therapies that rely on p53 activation. Indeed, re-expression of wild-type p53 by transient transfection in *TP53* knockout HCT116 cells restored the ability of Actinomycin D to induce transcription of canonical p53 targets p21 and MDM2. In contrast, expression of the p53 R175H mutant failed to reconstitute this response (**Fig. 3e**), confirming the functional impairment of this hotspot mutant in the context of IRBC activation. The R175H mutation, along with other structural hotspot non-sense mutations, accounts for approximately 25-40% of advanced CRC cases. Importantly, recent evidence indicates that the FDA approved Arsenic Trioxide (ATO) can restore the transcriptional activity of many p53 mutants, including the R175H [25]. We therefore hypothesized that pharmacological reactivation of mutant p53 could resensitize CRC cells to IRBC-inducing agents and to MLD regimens. *TP53* KO HCT116 cells expressing the R175H mutant were treated with increasing concentrations of ATO alone or in combination with Actinomycin D. In line with our hypothesis, co-administration of ATO restored the ability of mutant p53 to respond to Actinomycin D, leading to induction of p21 and MDM2 levels comparable to those observed with wild-type p53 (**Fig. 3e**). Based on this, *TP53* KO HCT116 cells and *TP53* KO cells reconstituted with p53 R175H were treated with ATO alone, AFOC (MLD) alone, or the combination of ATO + AFOC. While AFOC MLD had minimal impact in *TP53* KO cells and in R175H-expressing cells, co-administration of ATO restored sensitivity to the quadruple MLD regimen specifically in the R175H background (**Fig. 3f** and **S4b)**. Collectively, these findings demonstrate that pharmacological rescue of mutant p53 expands the therapeutic window of IRBC targeting and suggest that MLD regimens can be adapted to genetically advanced CRC by integrating p53-reactivating agents.

## Discussion

Ribosome biogenesis has long been considered an attractive therapeutic target in cancer, yet its pharmacological inhibition has been largely interpreted as a non-selective anti-anabolic intervention. Previous therapeutic efforts have primarily focused on maximizing suppression of rRNA synthesis, most notably through RNA polymerase I inhibitors such as CX-5461 [33]. Other experimental Pol I inhibitors, including BMH-21, have further established the vulnerability of cancer cells to disruption of ribosomal RNA synthesis but remain at preclinical stages. In these settings, antitumor activity has generally been linked to extensive disruption of ribosome production and the resulting loss of biosynthetic capacity, with nucleolar surveillance activated as part of a broader stress response. Our findings challenge this view by demonstrating that the cellular response to ribosome biogenesis perturbation is intrinsically structured, comprising a temporally and mechanistically distinct p53-dependent checkpoint response and a later global suppression of protein synthesis. The inability to disentangle these outputs has likely limited the selective therapeutic exploitation of nucleolar stress.

In this context, our data provide evidence that these two responses can be functionally and pharmacologically uncoupled. By distributing sub-threshold perturbations across multiple steps of the rRNA life cycle, we show that it is possible to sustain IRBC engagement while avoiding the overt inhibition of ribosome production and the global anabolic collapse associated with conventional high-dose treatment. This shifts the therapeutic paradigm from maximal inhibition of ribosome production to selective activation of nucleolar surveillance, reframing ribosome biogenesis as a checkpoint-regulated vulnerability rather than a purely metabolic dependency.

A key enabling element of this study is the use of a live-cell reporter for IRBC activation. While the molecular basis of the 5S-RNP–MDM2–p53 axis has been extensively characterized, its dynamic engagement in response to pharmacological perturbations has remained difficult to monitor. The identification of nucleolar L11 relocalization as a real-time and pathway-associated readout provides a direct link between drug action and checkpoint activation, and offers a practical framework for rational dose selection and combination design.

Our findings also provide a conceptual rationale for the use of multiple low-dose (MLD) strategies in this context. While MLD approaches have been successfully applied to inhibit oncogenic signaling pathways [21], here we extend this logic to the activation of a tumor suppressive checkpoint. By targeting distinct nodes of ribosome biogenesis at sub-effective doses, we achieve coordinated engagement of IRBC without triggering the broad biosynthetic suppression associated with high-dose monotherapies. The suppression of adaptive outgrowth observed under prolonged treatment suggests that sustained checkpoint activation may be sufficient to constrain tumor proliferation in a durable manner. At the same time, the present study primarily establishes a biological principle. Translation of a four-drug MLD regimen will require careful evaluation of dose scheduling, pharmacokinetic compatibility, cumulative toxicity and therapeutic windows in appropriate in vivo models. Although the compounds used here are either clinically approved or have entered clinical development, substantial preclinical work will be required to determine whether selective IRBC engagement can be achieved in vivo.

Finally, the restoration of IRBC responsiveness in cells harboring the structural p53 mutant R175H highlights the possibility to extend this principle beyond *TP53* wild-type tumors. Given the high prevalence of *TP53* mutations in advanced colorectal cancer, combining nucleolar stress–based approaches with mutant p53 reactivation provides a proof-of-concept for broadening checkpoint responsiveness, although its therapeutic applicability will require further validation.

Together, these findings establish a framework in which ribosome biogenesis perturbation can be tuned to preferentially activate nucleolar surveillance while limiting downstream anabolic collapse, supporting a model of “checkpoint-selective” therapeutic paradigm that may improve both efficacy and tolerability.

## Supporting information

Primers, Antibodies, Cells, Plasmids

## ACKNOWLEDGMENTS

We thank past and present members of the Cancer Metabolism Group, Dr. Lu who kindly shared the p53 R175H expression plasmid. This study was funded by Ministerio de Ciencia, Innovación y Universidades, (SAF2017-84301-P, PID2021-122125NB-I00, PID2025-172865NB-I00), the “Associación Española Contra el Cancer” AECC (LABAE20040GENT to A.G., CGB14142035THOM to R.S.) and by the Catalan Government (2017SGR01743, 2021SGR01026). The manuscript has been scanned using AI tools for typographical and minor editorial corrections.

## AUTHOR CONTRIBUTIONS

This study was conceived and directed by A.G., designed by A.G. and S.M.. S.M. and A.G. produced the Venus-L11 knock-in cellular model. S.M. carried out Venus-L11 immunofluorescence experiments, together with B.F., and performed IRBC complex immunoprecipitations. B.F. and A.G. carried out polysome profiling. AK and AKM tumor colon organoids experiments were executed by B.F.. Z.M. carried out RNA metabolic labeling, proliferation assays together with P.B., and reconstitution experiments with the p53 R175H mutant. *TP53* KO HCT116 and RKO isogenic cells lines were prepared by A.G, along with AK and AKM Colon organoids infections and selections. F.M. prepared the 3D spatial models of human 80S ribosomes. All authors analyzed the data and reviewed the manuscript. A.G. guided the studies and wrote the manuscript.

## Material and Methods

Polysome profiling, western blotting and Crispr targeting have been described in our previous publications [30, 34, 35]. All experimental details are provided in the Supporting Information (SI).

---

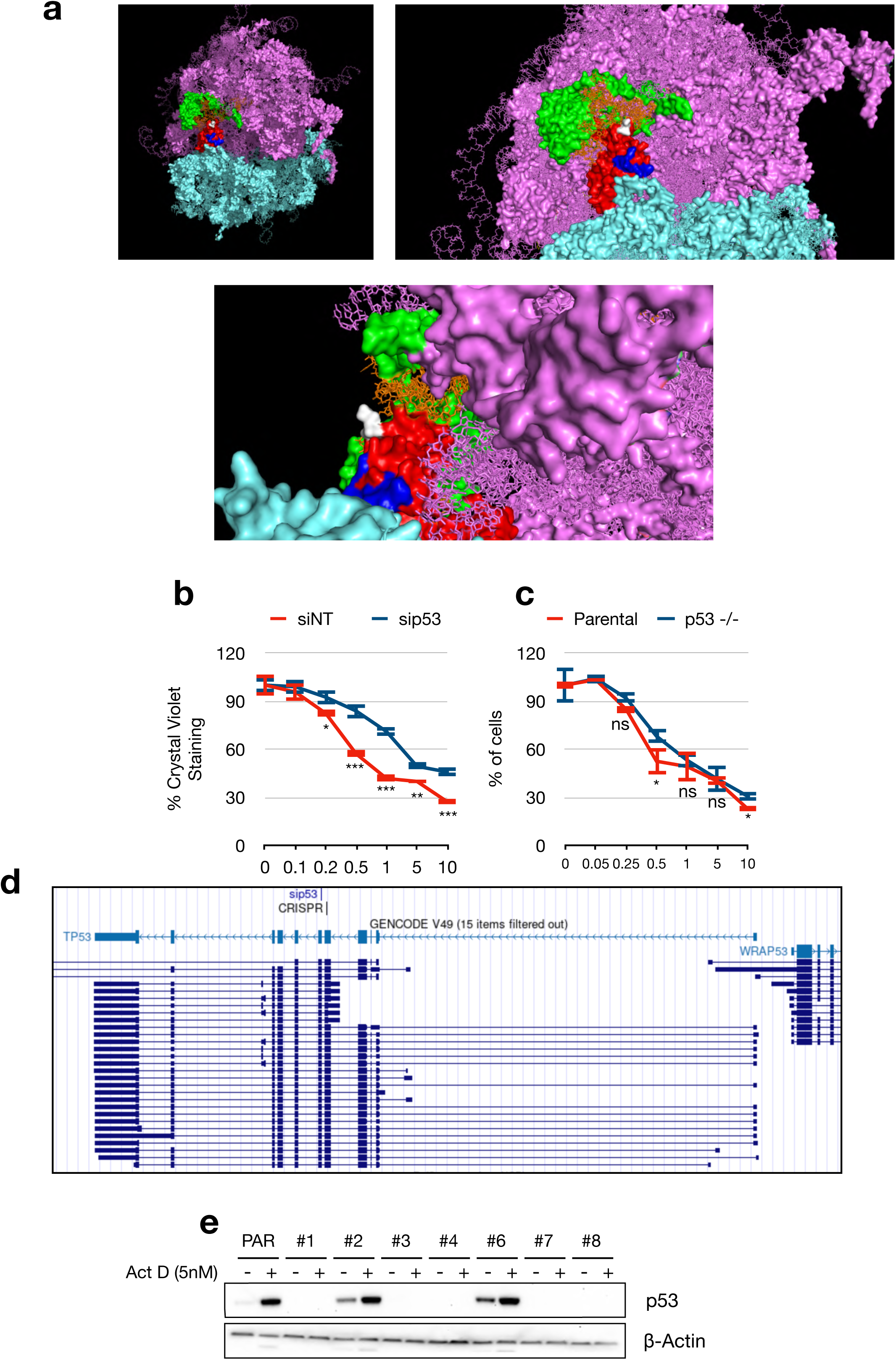

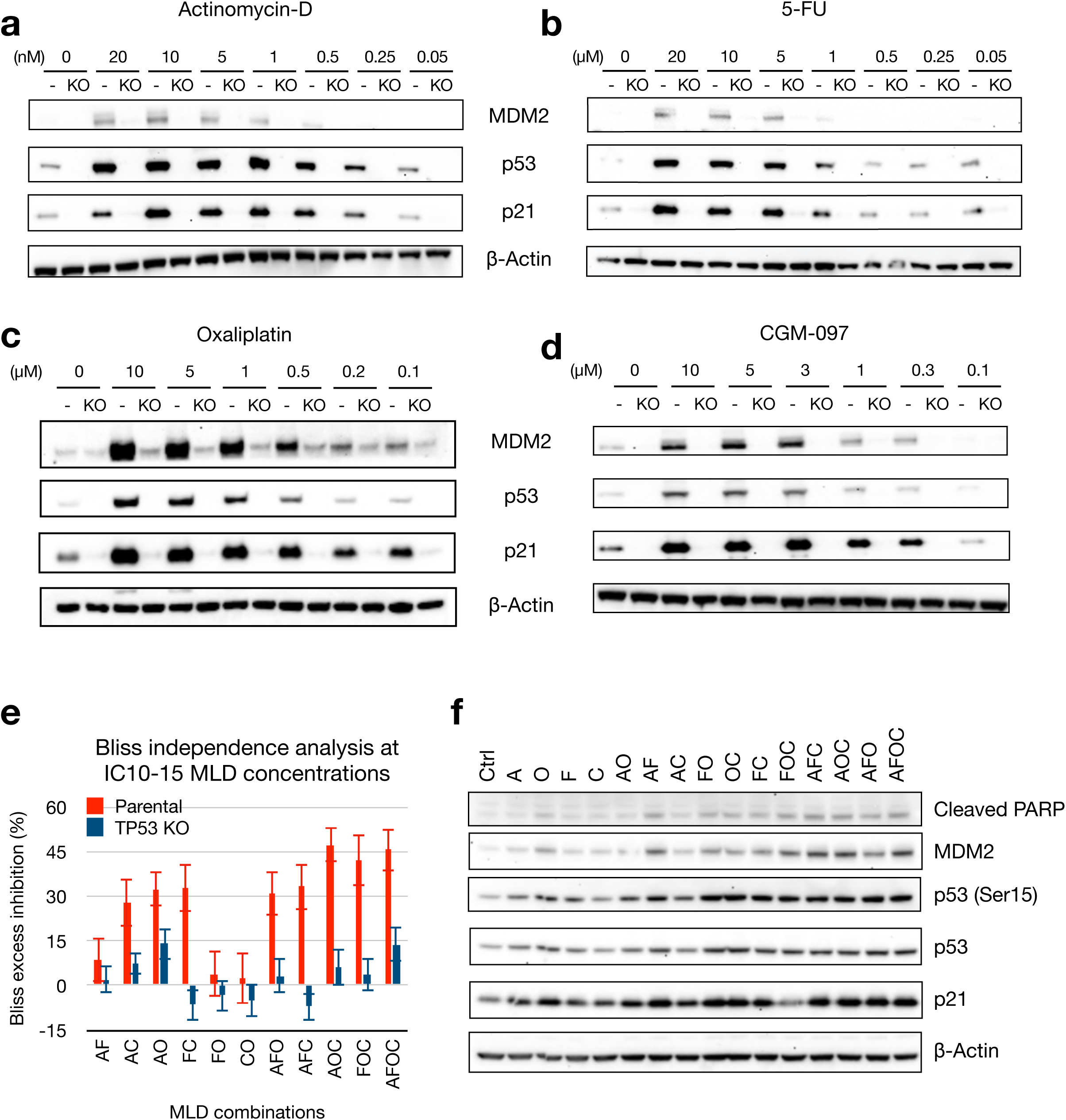

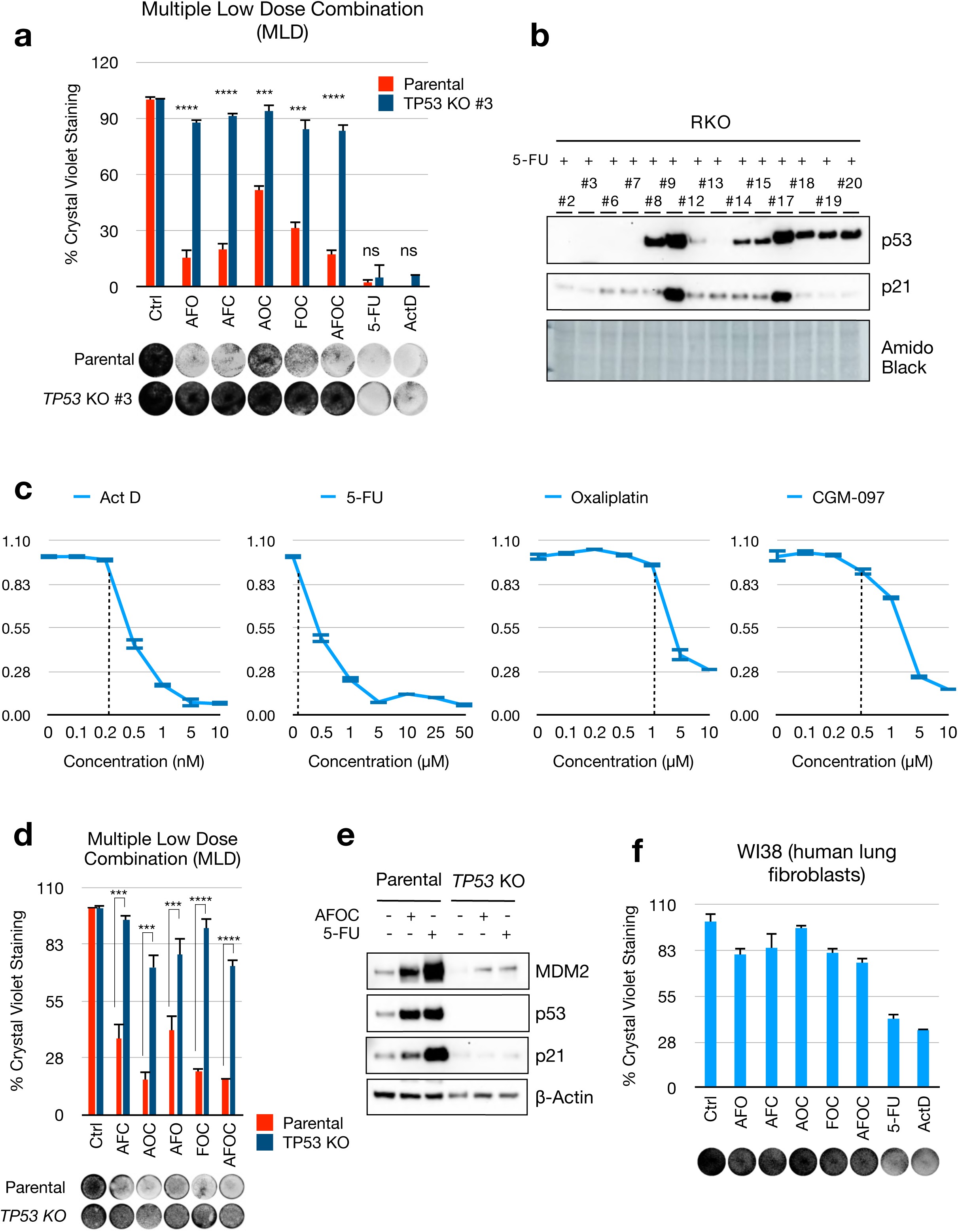

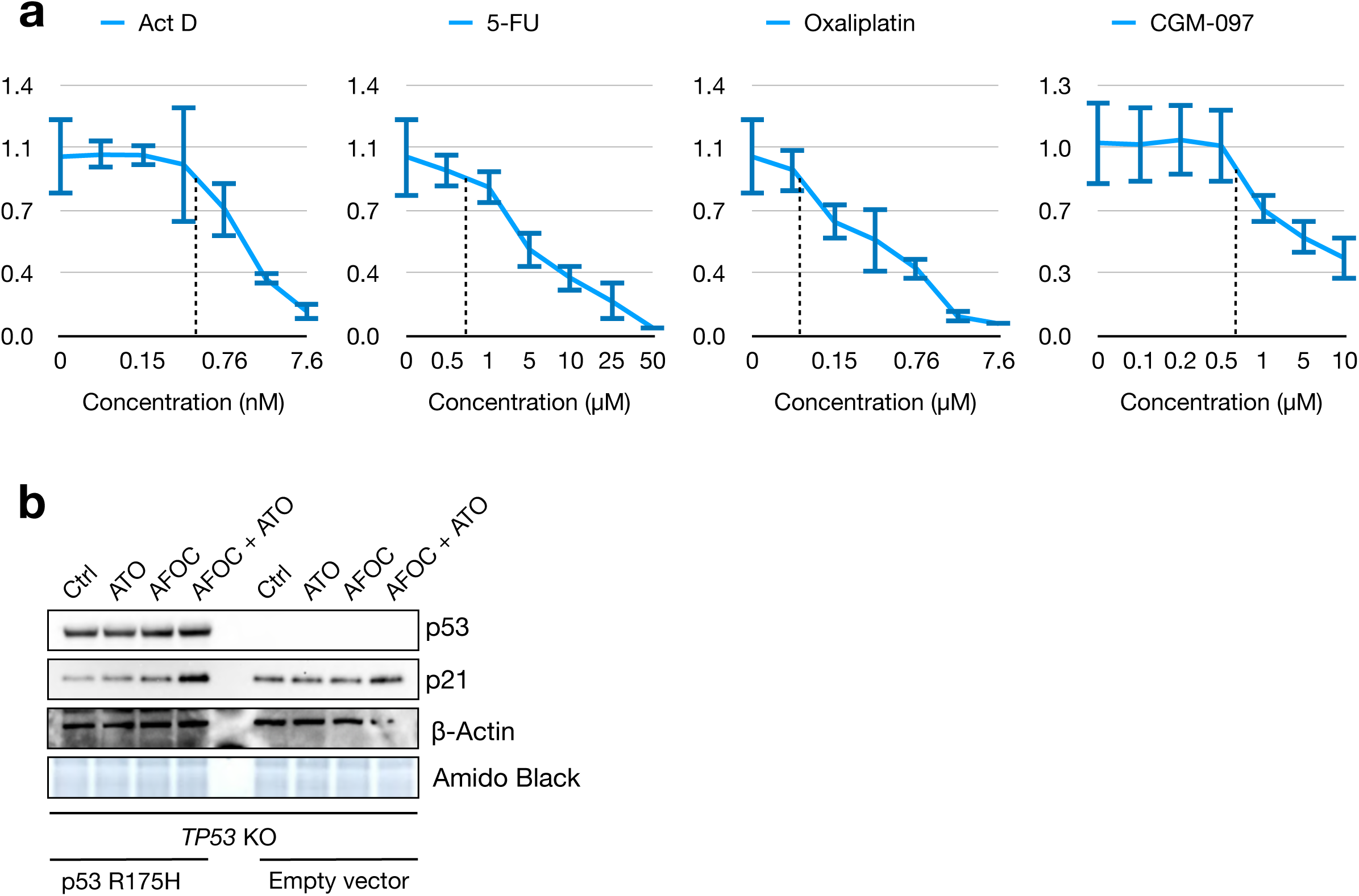

