## Supplementary material for "Pharmacologic decoupling of IRBC activation from anabolic collapse redefines ribosome biogenesis inhibition as a selective tumor suppressive strategy": Material and Methods, Supp. Figure Legends

### SUPPLEMENTAL INFORMATION

#### Supplementary Figure Legends

##### FIGURE S1 | Structural context of the IRBC complex and generation of TP53 knockout cells

(a) Structural representation of the human ribosome highlighting the IRBC complex components.
<u>Upper left panel</u>: overview of the 80S ribosome showing the spatial localization of L5 (green), L11 (red), and 5S rRNA (orange) within the large ribosomal subunit. The 40S subunit (18S rRNA and associated ribosomal proteins) is shown in light blue, whereas the 60S subunit (28S rRNA and associated ribosomal proteins) is shown in purple.
<u>Upper right panel</u>: magnified view of the IRBC region within the 60S subunit highlighting the L5–L11–5S rRNA complex.
<u>Bottom panel:</u> detailed view of L11 showing the N-terminal region (white) and C-terminal region (light blue) relative to the surrounding ribosomal environment.

(b) Crystal violet viability assay of HCT116 parental cells transfected with control siRNA (siNT) or siRNA targeting TP53 (sip53) following treatment with increasing concentrations of Actinomycin D.

(c) As shown in panel b, dose–response analysis comparing parental HCT116 cells and the previously reported HCT116 p53$^{-/-}$ cells generated by targeting exon 2 of *TP53*. Comparable sensitivity to Actinomycin D suggests residual IRBC responsiveness in this model.

(d) Genomic organization of *TP53* genomic locus and annotated transcript variants (GENCODE annotation) highlighting the location of exon 5 targeted for CRISPR–Cas9 editing, a region shared by all described TP53 transcript variants.

(e) Western blot screening of CRISPR-edited HCT116 clones in normal growing condition or following treatment with Actinomycin D (5 nM). p53 and β-Actin were measured.

##### FIGURE S2 | p53 pathway activation following ribosome biogenesis perturbation

(a) Western blot analysis of parental and *TP53* KO HCT116 cells treated with increasing concentrations of Actinomycin D, (b) 5-FU, (c) Oxaliplatin, or (d) the MDM2 inhibitor CGM-097. Stabilization of p53 and induction of canonical transcriptional targets (p21 and MDM2) in *TP53*-proficient and KO cells were detected. β-Actin served as loading control.

(e) Bliss independence analysis of the multiple low-dose drug combinations shown in Fig. 2c. Bliss excess inhibition was calculated from the residual viability values obtained for the corresponding single agents and drug combinations in Fig. 2c. For each combination, the expected residual viability under Bliss independence was determined as the product of the residual viabilities produced by the individual drugs, and Bliss excess was calculated as the difference between expected and observed residual viability. Positive values indicate greater growth inhibition than predicted under Bliss independence, values close to zero indicate independent effects, and negative values indicate lower-than-expected inhibition. Bars represent Bliss excess inhibition and error bars indicate propagated standard deviation calculated from the technical triplicates. The dashed line denotes Bliss independence (Bliss excess = 0).

(f) Immunoblot analysis of parental HCT116 cells treated with single agents or combinatorial multiple low-dose (MLD) regimens. Progressive engagement of the p53 pathway is observed with increasing combinatorial targeting, as indicated by p53 stabilization, p21 induction, phosphorylation of p53 (Ser15), MDM2 accumulation, and PARP cleavage.

**FIGURE S3 | Testing MLD regimens on RKO colorectal cancer cells**

(a) HCT116 parental cells and an independently generated *TP53* KO clones (#3) were treated with the indicated triple and quadruple multiple low-dose combinations or with standard-dose 5-FU (10 µM) and Actinomycin D (10 nM). Cell growth was assessed by Crystal Violet staining and expressed as percentage of the untreated control. Bars represent mean ± SD of triplicates. Representative images of Crystal Violet-stained wells are shown below the graph. A, Actinomycin D; F, 5-fluorouracil; O, oxaliplatin; C, CGM-097. Statistical comparisons were performed between *TP53* KO and parental cells within each treatment condition.

(b) RKO clones were generated by CRISPR-Cas9 targeting exon 5 of *TP53*. Clones were treated with 10 µM 5-FU for 24 hours and analyzed by western blot for p53 and p21 expression. Total protein staining is shown as loading control. Clone #6 was selected for subsequent experiments.

(c) Dose–response curves of parental RKO cells treated for 4 days with Actinomycin D, 5-FU, Oxaliplatin, or CGM-097 to define the RKO-specific IC10–15 concentrations used in subsequent MLD experiments. Cell viability was assessed by crystal violet staining from three biological replicates per condition, with untreated controls set as 100% viability. The dashed lines indicate the selected low-dose concentrations.

(d) Parental and *TP53* KO RKO cells (clone #6) were treated for 4 days with the indicated triple and quadruple MLD combinations. Cell growth was assessed by crystal violet staining and expressed relative to untreated controls. Bars represent mean ± SD from three biological replicates; representative stained wells are shown below. Statistical comparisons were performed between parental and TP53-knockout cells within each treatment condition.

(e) Parental and *TP53* KO RKO cells were treated for 24 hours with AFOC or 10 µM 5-FU. Whole-cell lysates were analyzed by western blot for MDM2, p53 and p21 expression. β-Actin was used as loading control.

(f) WI38 human embryonic lung fibroblasts were treated for 4 days with the indicated triple and quadruple MLD combinations or with standard-dose 5-FU (10 µM) and Actinomycin D (10 nM). Cell growth was assessed by crystal violet staining and expressed relative to untreated controls. Representative stained wells are shown below the graph.

**FIGURE S4**

(a) Dose–response curves of AK mouse tumor colon organoids treated for 4 days with Actinomycin D, 5-FU, Oxaliplatin, or CGM-097 to define the IC10–15 concentrations used in subsequent MLD experiments. Cell viability was measured using CellTiter-Glo and normalized to untreated controls. Data are shown as mean ± SD from six replicate wells per condition. Dashed lines indicate the selected low-dose concentrations.

(b) Western blot analysis of HCT116 *TP53* KO cell lysates from Figure 3f.

**Supplementary Material and Methods**

**Cell culture and reagents**

HCT116 and RKO colorectal cancer cell lines, and WI38 human embryonic lung fibroblasts were obtained from the ATCC and maintained in DMEM (Thermo Fisher Scientific) supplemented with 10% heat-inactivated FBS (Sigma-Aldrich). Transfections of siRNAs, were carried out using RNAiMax (Thermo Scientific) at a concentration of siRNAs of 20nM. For all studies, cells were incubated at 37°C, 5% CO2, and 90% to 95% of relative humidity. Actinomycin-D was from SelleckChem, 5-FU and ATO from SIGMA, Oxaliplatin and CGM-097 from MedChemExpress. The BCA Reagents were from Pierce. The primary antibodies and their corresponding working dilutions utilized in this study are listed in Supplementary Table S1. MagnaCHIP Protein A/G magnetic beads mix were from Millipore. Goat anti-(mouse IgG)-peroxidase conjugate and goat anti-(rabbit IgG)-peroxidase conjugate were from Dako. The Placental Rnase inhibitor was purchased from NEB (New England Biolabs). The sequences of siRNAs used in the experiments, siNS, siRPS6 and siRPL7a are reported in Supplementary Table S1.

**Mouse strains**

Apc$^{fl/fl}$ mice carrying loxP sites flanking exon 14 of *Apc* and *Kras*$^{LSL\text{-}G12D/+}$ mice were obtained from The Jackson Laboratory [1, 2]. The Mdm2$^{C305F/C305F}$ strain was kindly provided by Dr. Zhang [3]. Mouse lines were intercrossed to generate animals carrying the following genotypes: *Apc*$^{fl/fl}$;*Kras*$^{LSL\text{-}G12D/+}$ (AK) and *Apc*$^{fl/fl}$;*Kras*$^{LSL\text{-}G12D/+}$;*Mdm2*$^{C305F/C305F}$ (AKM). Genotypes were confirmed by PCR-based genotyping using primers specific for the respective alleles.

**Cell Proliferation Assay**

Cell proliferation was assessed using a crystal violet staining assay. Cells were seeded in 12-well plates at a density of 50,000 cells per well in triplicate for each experimental condition. Treatments were administered and at the indicated time points (4 days or 30 days) cells were fixed at room temperature (RT) with 4% paraformaldehyde for 10 minutes. Following fixation, cells were stained with a crystal violet solution (0.5% crystal violet and 10% methanol in distilled water) for 15 minutes at RT. Plates were then thoroughly washed with phosphate-buffered saline (PBS) to remove excess dye and allowed to air dry. Images of the stained wells were captured and analyzed using ImageJ software and densitometric analysis was carried out using non treated control as 100% of viability. For 30 days treatments drugs combinations were refreshed every 4 days.

**Generation of TP53 Knock-out and Venus-RPL11 Knock-in cells by CRISPR-Cas9**

To knockout *TP53* gene HCT116 and RKO cells were transfected with pX458 plasmid expressing a guide strand (sgRNA) sequence targeting exon 5 of the TP53 gene using Lipofectamine 2000 transfection reagent (Invitrogen). As the pX458 plasmid contains GFP, single-cell sorting was performed 48 hours after transfection using a MOFLO Astrios (BD Biosciences), and individual clones were cultured and subsequently analyzed for p53 protein expression by immunoblotting, leading to the isolation of the *TP53 knock-out* clones (KO#8 and #3 for HCT116 and KO#6 for RKO). Primers targeting the exon 5 of *TP53* gene are reported in supplementary Table 1. To knock-in the translational start site of *RPL11* with Venus fluorescent protein coding sequence, a guide strand targeting the exon 1 of *RPL11* was cloned in pX330 and a targeting vector including a ~500 bp homology arms upstream and downstream of the sgRNA targeting site and separated by the Venus CDS in frame with the L11 CDS was cloned into pL253 (L11 targeting-pL253) and co-transfected in HCT116 cells. 48 hours from transfection the Venus-positive cells, deriving from a bona fide homology directed repair in frame with RPL11 coding sequence, were sorted as single cells and monoculture were expanded and characterized by western blot with anti-RPL11 and anti-GFP antibodies as well as by confocal microscopy.

**Protein analysis**

Cell protein extracts for Western blot analysis were prepared by using a 1% SDS lysis buffer [50 mM tris (pH 7.4) and 1% SDS] supplemented with the protease inhibitor cocktail (Sigma-Aldrich), phosphatase inhibitor cocktail 2 and 3 (MedChemExpress), and benzonase (100 U/ml, Millipore). After lysis, cell lysates were incubated 30′ on ice followed by sonication and centrifugation at 13,000 rpm for 10′. Alternatively, cell pellets were lysed in RIPA buffer as previously described [4]. Protein concentrations were determined for supernatants by the BCA assay (Pierce). Protein extracts (20-30 µg) were resuspended in Laemmli SDS sample buffer (1×) and boiled at 95°C for 10 min. Proteins were separated on 10% SDS–polyacrylamide gels by electrophoresis and transferred to polyvinylidene difluoride (PVDF) membranes (GE Healthcare Life Science). Blots were stained with amido black to confirm equal loading and transfer of proteins, blocked with 5% non-fat dry milk and then probed with the indicated antibodies. Immunoblots were developed using secondary horseradish peroxidase–coupled antibodies and an enhanced chemiluminescence kit (GE Healthcare).

**Polysome profile analysis**

Distribution of mRNAs across sucrose gradients was performed as described earlier [5], except for minor modifications. Briefly, cells were plated in 150-mm dish and subjected to the indicated treatment. Cycloheximide (CHX) was added to the medium at 37°C for 5 min at a concentration of 100 µg/ml. Cells were washed twice with cold PBS supplemented with CHX, scraped on ice, and pelleted by centrifugation at 3000 rpm for 3′ in cold. Cell pellets were resuspended in 250 µl of hypotonic lysis buffer [1.5 mM KCl, 2.5 mM MgCl2, 5 mM tris HCl (pH 7.4), 1 mM dithiothreitol (DTT), 1% sodium deoxycholate, 1% Triton X-100, and CHX (100 µg/ml)] supplemented with mammalian protease inhibitors (Sigma-Aldrich) and RNase inhibitor (NEB) at a concentration of 100 U/ml and left in ice for 5′. Cell lysates were cleared of debris and nuclei by centrifugation for 5′ at 13,000 rpm. Protein concentrations were determined by BCA assay, and equal amount of polysomal lysate (500 to 1000 µg, depending on the experiment) was loaded on 10 to 50% sucrose linear gradients generated with a BIOCOMP gradient master and containing 80 mM NaCl, 5 mM MgCl2, 20 mM tris HCl (pH 7.4), 1 mM DTT, and RNase inhibitor (10 U/ml). Gradients were centrifuged on a SW40 rotor for 2 hours and 30 min at 35,000 rpm. Gradients were analyzed on a BIOCOMP gradient station detecting both OD 260 nm (rRNA) and 480 nm (Venus-L11) and collected in 12 or 13 fractions ranging from light to heavy sucrose. Fractions were supplemented with SDS at a final concentration of 1% and placed for 10 min at 65°C [6]. Venus-RPL11 distribution on sucrose gradients was analyzed by standard TCA precipitation as described earlier [7]. Briefly twenty percent volume of each fraction was utilized for protein precipitation, which was resuspended in 1X Laemmli SDS sample buffer and boiled for 10 min at 95°C, resolved by SDS-

PAGE and transferred to a PVDF membrane. The membrane was stained with Ponceau Red and incubated with the corresponding antibodies.

**Immunoprecipitations**

Cells grown in 15 cm dishes were lysed on ice in immunoprecipitation lysis buffer [50 mM Tris HCl (pH 7.5), 150 mM KCl, 5 mM MgCl2, 1 mM EGTA, 1mM DTT, 10% glycerol, 0.8% NP-40, PMSF 1mM, 100 U/ml RNaseout inhibitor (Invitrogen) and complete, EDTA-free Protease Inhibitor cocktail (Roche)] and subjected to immunoprecipitation, largely as previously described [8]. In brief, ribosomes were pelleted by ultracentrifugation at 200,000 g at 4°C for 2 h. Immunoprecipitations were obtained from equivalent amounts of protein incubated with 20 µl of GFP-TRAP beads (Proteintech) at 4°C overnight rotating. Supernatants were discarded and beads washed three times and resuspended in protein loading buffer for Western blot analysis.

**Immunofluorescence**

Cells were grown in 2-well glass chamber slides. Cells were fixed with 4% (w/v) PFA for 10 min at room temperature. After a 0.2% Triton X-100 permeabilization step for 5 min at room temperature, non-specific antibody binding sites were blocked with 10% (v/v) normal goat serum. After an overnight incubation of cells with a 1:400 dilution of monoclonal anti-p53 antibody in 2% normal goat serum, cells were incubated with Anti-Mouse Alexa Fluor 555 1:800. Coverslips were mounted

using Vectashield with DAPI (Vector Labs, Burlingame, CA). Fluorescence was detected with the Leica spectral confocal microscope TCS SP5 using a 63X N.A 1.4 objective and LAS AF software. Fluorophores were excited with Argon laser for 488 nm, DPSS 561 for 555 nm and Diode laser for 405 nm. Images were analyzed with FIJI software (NIH). Venus-L11 nuclear/nucleolar accumulation was quantified by calculating the nuclear/cytoplasmic ratio (Fig. 1h and 2d).

For real time live-cell imaging, HCT116 L11-Venus cells transiently expressing mPlum-FBL were grown onto glass bottom 8-well slides (IBIDI). Live-cell imaging was performed on the Leica spectral confocal microscope TCS SP5. Images were taken every 10 minutes for a total time of 4 hours using the 63x glycerol objective.

**Detection of Nascent rRNA Synthesis**

For metabolic labeling of rRNA species by Northern blot analysis, cells were treated as indicated. After 24 hours nascent RNA was labeled by adding to the medium 5-Ethynyl-Uridine (EU) at a concentration of 200 µM for additional 2 hours, then cells were harvested. Total RNA was extracted with TRIZOL and 3ug of total RNA were resolved by Northern Blot. Nascent RNA was detected by Biotin-Azide click reaction on the membrane, streptavidin-HRP incubation and ECL detection. Ethidium Bromide staining of the gel detected RNA integrity. Cells not pulsed with EU served as negative control for metabolic labeling.

**Tumor Colon Organoids preparation and analysis**

Colon organoids were prepared from animals with the following genotypes: $Apc^{fl/fl}$; $Kras^{LSL\text{-}G12D/+}$ (AK) and $Apc^{fl/fl}$; $Kras^{LSL\text{-}G12D/+}$; $Mdm2^{C305F/C305F}$ (AKM) [1-3]. Colon crypts were isolated as previously described [9, 10]. Isolated crypts were then mixed with growth factor-reduced Matrigel

(Corning) at a 1:4 ratio (crypt suspension:Matrigel) and fully supplemented Intesticult™ medium (Stem Cell Technologies) was added to each well. Crypts were cultured to allow the formation of typical colon organoids structures. Organoids were then mechanically dissociated and transduced with a Cre-expressing adenovirus for 4 h at 37°C to induce *Apc* deletion and activation of the *Kras*$^{G12D}$ allele. Following infection, organoids were re-embedded in Matrigel and maintained in IntestiCult™ medium. Five days later, the medium was replaced with a minimal selection medium consisting of Advanced DMEM/F-12 supplemented with N2 and B27, thereby selecting for niche-independent tumor colon organoids. Established AK and AKM tumor organoids were expanded, cryopreserved and routinely genotyped to confirm Cre-mediated recombination. For drug-response assays, established AK and AKM tumor colon organoids were dissociated to single cells and 1,000 cells per well were seeded in 96-well plates in 10 $\mu$L Matrigel-containing domes. Six technical replicates were used for each condition. Organoids were cultured in minimal selection medium alone or supplemented with the AFOC combination, using concentrations defined from IC10 dose-response analyses performed in AK tumor colon organoids, or with a standard high dose of 5-FU (10 $\mu$M). After 4 days of treatment, cell viability was measured using CellTiter-Glo (Promega) according to the manufacturer's instructions.

**Puromycin Incorporation Assay**

Global protein synthetic capacity was assessed by puromycin incorporation, which relies on the incorporation of puromycin—an aminonucleoside antibiotic that mimics aminoacyl-tRNAs—into nascent polypeptide chains, allowing their detection by immunoblotting. Cells were treated as indicated for 24 hours then incubated with puromycin (Sigma-Aldrich) at a final concentration of 10 µg/mL for 30 minutes at 37 °C. After incubation, cells were washed with ice-cold PBS, lysed, and processed for Western blot analysis using an anti-puromycin antibody (clone 12D10, Millipore). Signal intensities per lane generated by anti-puromycin antibody and Amido black were measured by ImageJ.

**Bliss independence analysis**

Bliss independence analysis was performed at the predefined low-dose concentrations used in the MLD regimens. Viability values were normalized to untreated controls and expressed as fractional residual viability. For each combination, the expected residual viability was calculated as the product of the residual viabilities produced by the corresponding single agents. Bliss excess inhibition was defined as the difference between expected and observed residual viability, with positive values indicating greater inhibition than predicted under Bliss independence. Means and standard deviations were calculated from technical triplicates. Errors associated with the expected values were propagated from the corresponding single-agent measurements, and the standard deviation of the Bliss excess was calculated by combining the propagated expected-value error with the standard deviation of the observed combination response. Single-agent and combination measurements obtained within the same experiment were used whenever available.

**Statistical analysis**

Statistical analyses were performed using GraphPad Prism version 8.0. Data are presented as mean ± SD unless otherwise indicated. Comparisons between two experimental groups were performed using two-tailed unpaired Student's t-test. Statistical significance was defined as $p < 0.05$ ($p < 0.05$, *$p < 0.01$, **$p < 0.001$, ***$p < 0.0001$).
