## Supplementary material for "Pharmacologic decoupling of IRBC activation from anabolic collapse redefines ribosome biogenesis inhibition as a selective tumor suppressive strategy": Primers, Antibodies, Cells, Plasmids

| primer name | Sequence 5'-3' | Use | |
|---|---|---|---|
| LeftARM_F_BamHI | ACGGATCCCCTTCAGACTATGTGTATAAGGTG | Left Arm cloning of Venus-L11 targeting vector | |
| Left ARM-R-new (BsaI) | ACGGTCTCGATGGAGAGCAGGAAGAGA | Left Arm cloning of Venus-L11 targeting vector | h.s. = homo sapiens |
| Venus-F-new (BsaI) | ACGGTCTCTCCATCATGGTGAGCAAGGGCGAG | Venus cloning of Venus-L11 targeting vector | |
| Venus_R_BsaI | ACGGTCTCACCGCTCGAGATCTGAGTCCGGAC | Venus cloning of Venus-L11 targeting vector | |
| RightARM_F_BsaI | ACGGTCTCAGCGGTGAGTAGCTGGGACCTG | Right Arm cloning of Venus-L11 targeting vector | |
| RightARM_R_SalI | ACGTCGACCAAGAAACAACCAAGCGACC | Right Arm cloning of Venus-L11 targeting vector | |
| L11-Venus-CRI1_F | CACCGCCAGCTACTCACCGCCATGA | Cloning sgRNA targeting L11 exon 1 in pX330 | m.m. = mus musculus |
| L11-Venus-CRI1_R | AAACTCATGGCGGTGAGTAGCTGGC | Cloning sgRNA targeting L11 exon 1 in pX330 | |
| TP53_CR_EX5_F | CACCGCAGTCACAGCACATGACGG | Cloning sgRNA targeting TP53 exon 5 in pX458 | |
| TP53_CR_EX5_R | AAACCCGTCATGTGCTGTGACTGC | Cloning sgRNA targeting TP53 exon 5 in pX458 | |
| siNT | GCAUCAGUGUCACGUAAUA | siRNA | |
| siRPL5 | GCUUGGUGAUACAAGAUAA | siRNA | |
| siTP53 | GCAUCUUAUCCGAGUGGAA | siRNA | |

Antibodies

| Antibody | Brand | Catalog number | Working dilution |
|---|---|---|---|
| anti-p53 | Santa Cruz Biotechnology | sc-126 | 1:1000 |
| anti-p21 | BD Pharmingen | 554228 | 1:1000 |
| anti-HDM2 | Santa Cruz Biotechnology | sc-965 | 1:1000 |
| anti-GFP | Abcam | ab290 | 1:2500 |
| anti-L11 | Invitrogen | 37-3000 | 1:1000 |
| anti-L11 | Made by Dr. Sinisa Volarević | Clone H3 | 1:1000 |
| anti-phospho-p53 (Ser15) | Cell Signaling Technology | 9284 | 1:1000 |
| anti-β-Actin | SIGMA | A2228 | 1:10000 |
| anti-PARP1 | BD Biosciences | 51-6639GF | 1:1000 |
| anti-RPL5 | Bethyl | A303-933A | 1:2000 |

Cell Lines

| Cell line | Acronym | Origin | Reference |
|---|---|---|---|
| HCT116 | | ATCC | CCL-247 |
| HCT116 L11-Venus (T18) | | This study | CRL-3216 |
| HCT116 L11-Venus (T21) | | This study | CCL-75 |
| HCT116 TP53 KO | | This study | CRL-2577 |

Plasmids

| Plasmid name | Insert | Origin / Reference |
|---|---|---|
| p53 WT / pcDNA 3.1 | p53 WT CDS | Received from Dr. Lu, Chen et al., Cancer Cell 21 |
| p53 R175H / pcDNA 3.1 | p53 R175H CDS | Received from Dr. Lu, Chen et al., Cancer Cell 21 |
| L11 sgRNA / pX330 | L11 sgRNA targeting exon 1 | This study |
| p53 Ex5 sgRNA / pX330 | p53 sgRNA targeting exon 5 | This study |
| L11-Venus targeting vector / pL253 | L11-Venus targeting vector with homology arms for knocking | This study |
| mPlum-FBL | mPlum-Fibrillarin CDS | Addgene (Plasmid #55969) |
| pL253 | Empty vector | Liu et al., Genome Res. 2003 |
| pX330 | Cas9 and sgRNA | Addgene (Plasmid #42230) |
| pX458 | Cas9-GFP and sgRNA | Addgene (Plasmid #48138) |
